# The role of the thalamus in human reinforcement learning

**DOI:** 10.1101/2022.11.23.517731

**Authors:** Antoine Collomb-Clerc, Maëlle C. M. Gueguen, Minotti Lorella, Kahane Philippe, Navarro Vincent, Bartolomei Fabrice, Carron Romain, Regis Jean, Chabardès Stephan, Stefano Palminteri, Julien Bastin

## Abstract

Although the thalamus is supposed to be involved in reinforcement-based decision-making, there is no direct evidence regarding the involvement of this subcortical structure in humans. To fill this gap, we leveraged rare intra-thalamic electrophysiological recordings in patients and found that temporally structured thalamic oscillations encode key learning signals. Our findings also provide neural insight into the computational mechanisms of action inhibition in punishment avoidance learning.

## Main Text

As the philosopher, John Locke would put it “reward and punishment are the only motives to a rational creature: these are the spur and the reins whereby all mankind is set on work and guided”. Research in reinforcement learning aims at characterizing the processes through which people learn, by trial and error, to select actions that respectively maximize or minimize the occurrence of rewards or punishments^1^. Converging evidence suggests that reward-based reinforcement learning engages a fronto-striatal circuit and the dopaminergic system^2,3,4^. However, there is no evidence in humans regarding how neural activity in the thalamus - a key node in this circuit - encodes variables related to reinforcement learning processes.

Punishment avoidance learning is of equal ecological importance for organism survival and has been shown in many experimental settings to be at least as effective as reward seeking^5,6^. Critically, while the performance based on rewards or punishments exhibits comparable learning accuracies, subjects are constantly slower in punishment avoidance learning tasks^7^. This increase in reaction time is thought to reflect a manifestation of a Pavlovian bias according to which motor responses are inhibited by punishment expectations, irrespective of the appropriateness of the instrumental response^8,9,10^.

Intriguingly, this behavioral asymmetry between reward-seeking and punishment avoidance is mirrored by a neural asymmetry: the ventral striatum and ventromedial prefrontal cortex represent reward learning signals, while the amygdala, anterior insula, or lateral orbitofrontal cortex rather represent punishment learning signals^11,12,13,14^. Despite early lesion studies in rabbits^15^ suggesting the involvement of the mediodorsal and the anterior parts of the thalamus during punishment-avoidance learning, most of the animal studies in mice^16,17^, rats^18^, rabbits^19^, or monkeys^20,21^ surprisingly focused on reward-based learning, leaving the role of theses thalamic regions in punishment-based learning largely unexplored.

The high spatiotemporal resolution necessary to disentangle human thalamic neuronal activities during such cognitive processes is unattainable with ordinary imaging tools. Thus, we preferentially leveraged rare direct neural recordings in the human limbic thalamus. We investigated whether neuronal oscillations were associated with reinforcement-related signals at different time points during a well-validated reward-seeking and punishment avoidance learning task^5,11,12^. This combination of intra-thalamic recordings with computational modeling of the learning behavior results in the first time-resolved investigation of choice and learning signals in the human thalamus.

Local field potentials were recorded from eight drug-resistant epileptic patients (Table S1) implanted bilaterally in the thalamus with deep-brain stimulation electrodes as a surgical treatment to alleviate their seizures. Electrodes had two upper contact pairs inside the anterior thalamic nucleus, with the more ventral contact pairs localized in the dorsomedial thalamic nucleus (Fig. 1a). Intra-thalamic recordings were collected while patients were performing a previously validated instrumental learning task with the instruction to maximize the monetary gains and minimize the monetary losses (Fig. 1b)^5,11,12^.

**Figure 1.**
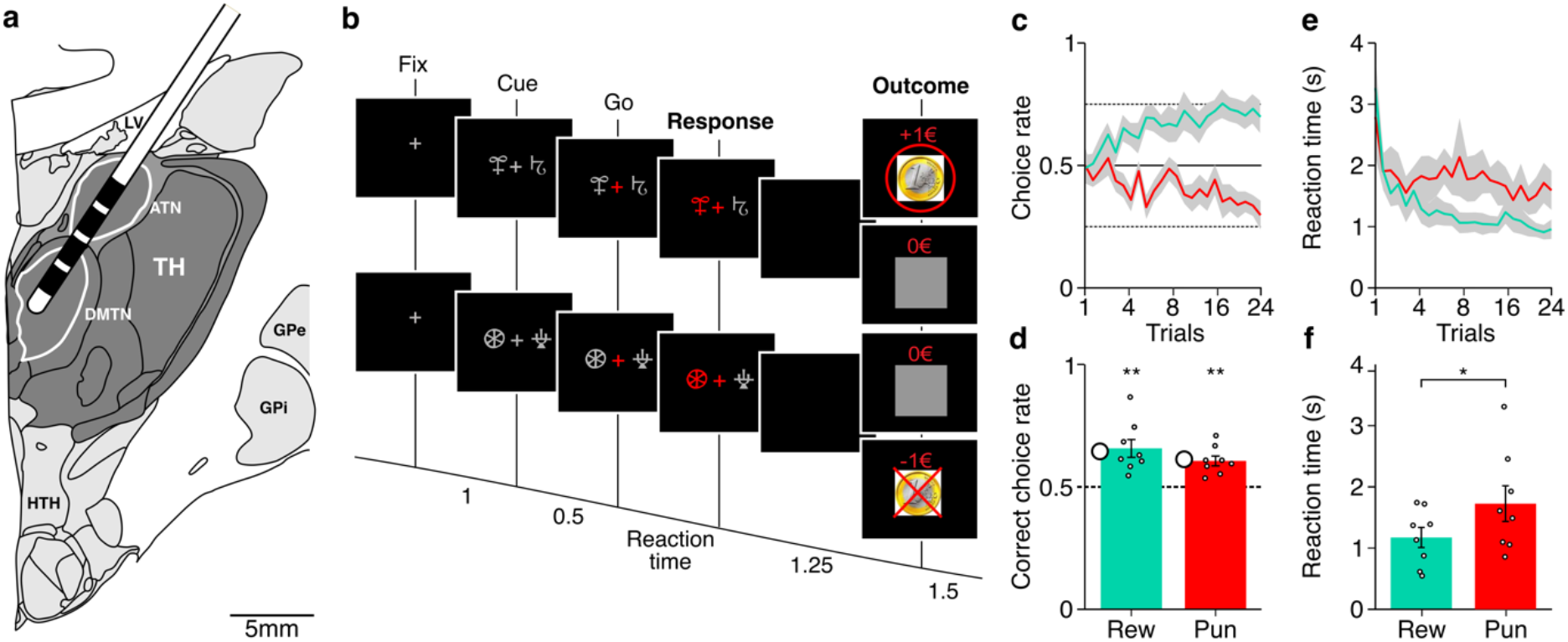
Reinforcement-learning paradigm and behavior. **a**. Schematic figure of the position of the deep brain stimulation electrodes used to record intra-thalamic signals (ATN: anterior thalamic nucleus; DMTN: dorsomedial thalamic nucleus; TH: Thalamus; HTH: Hypothalamus; GPi/GPe: Globus pallidus intern/extern; LV: Left ventricle). **b**. Successive screenshots of a typical trial in the reward (top) and punishment (bottom) conditions. Patients had to select one abstract visual cue among the two presented on each side of a central visual fixation cross and subsequently observed the outcome. Durations are given in seconds. **c**. Average±SEM learning curves across patients (n = 8) through trials shown separately for the reward (green) and punishment (red) conditions. **d**. Average±SEM choice performance across patients in the reward (Rew) and punishment (Pun) conditions. The average predicted performance from a fitted Q-learning model is indicated by a circle for each condition. Dots represent data from individual patients. Asterisk indicates the significance of the one-sample t-test used to compare for each condition the correct choice rate to the chance level (i.e., 50%). **e**. Average±SEM reaction times across patients (n = 8) through trials shown separately for the reward (green) and punishment (red) conditions **f**. Average±SEM reaction times across patients in the reward (Rew) and punishment (Pun) conditions. Dots represent data from individual patients. Asterisk indicates the significance of a paired t-test comparing reaction times between conditions.

Behavioral results were consistent with what was previously observed in this task (Fig. 1c-d). Accuracy was higher than chance in both the reward (65±0.04, t(7) = 4.23, p = 0.0039) and punishment conditions (0.60±0.02, t(7) = 5.13, p = 0.0014) and was not different between the two conditions (t(7) = 1.68, p = 0.14). Reaction times were significantly shorter in the reward (1173±164 ms) than in the punishment (1726±291 ms) condition (t(7) = −3.10, p = 0.017). Thus, patients learned similarly from rewards and punishments but took longer to choose between cues for punishment avoidance, in line with previous behavioral data from healthy subjects^7^ or epileptic patients^12^. These results confirm that, although instrumental performances are similar, the decision process differs in reward-seeking and punishment-avoidance contexts in a way that is compatible with a motor inhibition induced by punishment expectation^8,9,10^.

We next investigated the association between thalamic neural activity and reinforcement learning variables. We fitted a Q-learning model to the behavioral data of each patient to estimate trial-wise values of the expectation. The neural activity of each recording site (n=48 sites) was then regressed in the time-frequency domain against both expectation and outcome signals at different time points during the task. Given the absence of significant differences between sites located within the anterior thalamic nucleus (n=16 sites), the dorsomedial thalamic nucleus (n=16 sites, Supplementary. Fig. S1) or sites localized in-between (n=16), in the following, all the analyses were conducted across all recording sites.

We first investigated neural signals occurring after the cue (Fig. 2a) and before the choice onset (Fig. 2b). We found that low-frequency oscillations (LFOs, 4-12 Hz) were significantly correlated with punishment expectations (Qp) early after the cue onset (Fig. 2c; 0.36 to 1.14 s window, β_Qp_ = 0.33±0.02, sum(t(47)) = −36.38, p_c_<0.05) whereas there was no significant association between thalamic LFOs and reward expectation (Qr) at these latencies. Furthermore, we found that LFOs were associated more strongly with Qp than with Qr (Fig. 2c; 0.52 to 0.98s window, β_Qp_-β_Qr_ = 0.34±0.02, sum(t(47)) = 20.07, p_c_ < 0.05). Conversely, when neural activity was time-locked to the choice onset (Fig. 2b), there was a significant association between thalamic LFOs and expectations signals during both learning conditions (Fig. 2d; −2.22 to −0.81 s window, β_Qr_ = 0.21±0.01, sum(t(47)) = 75.00, p_c_ < 0.05; −1.44 to 0.03 s window, β_Qp_ = 0.35±0.02, sum(t(47)) = 75.24, p_c_ < 0.05). Altogether, decision-related activities in the thalamus are consistent with a stronger encoding of punishment expectations (Qp), at least during the first second after stimulus onset, although both reward and punishment expectations are encoded later on.

**Figure 2.**
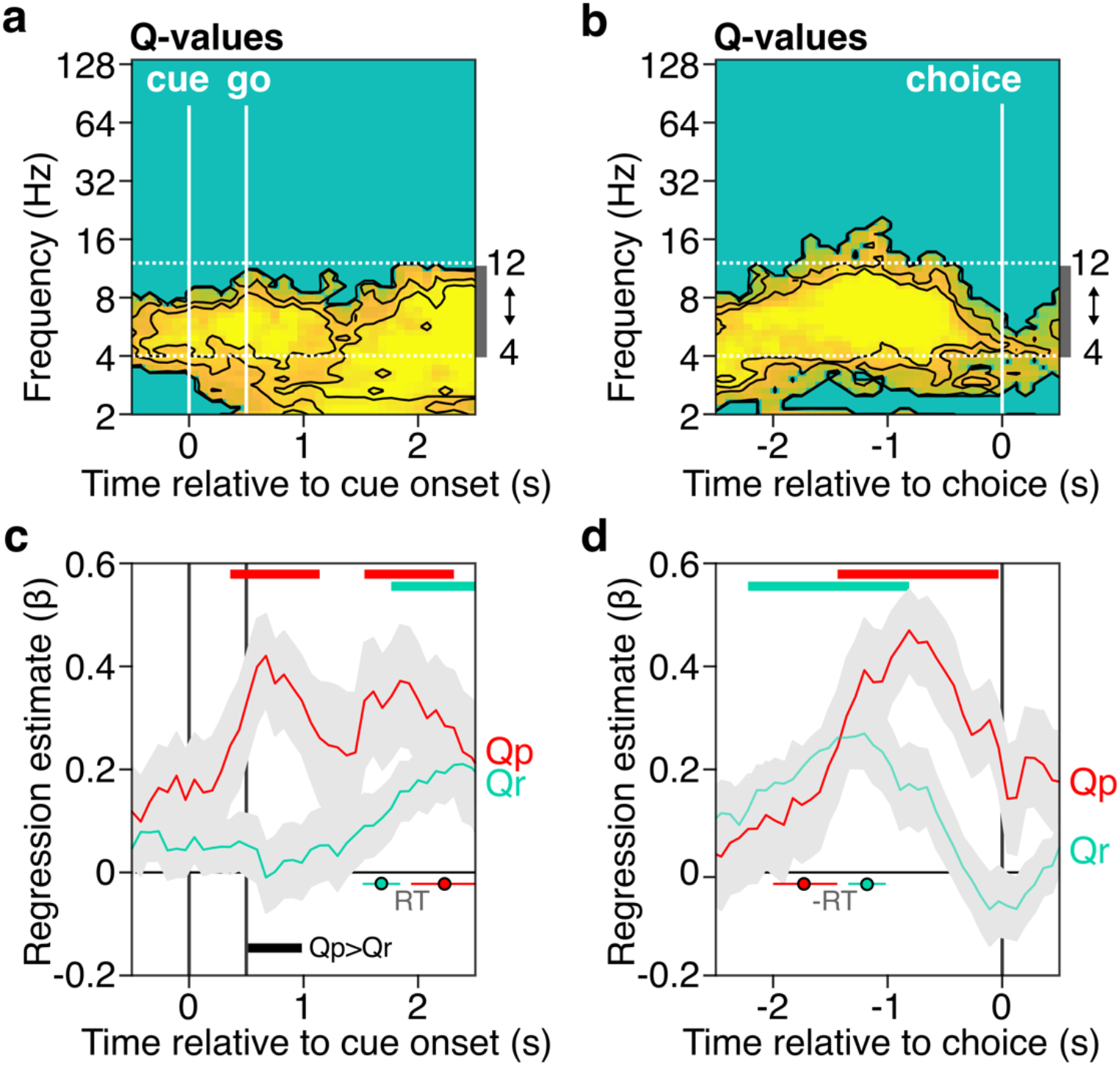
Thalamic low-frequency oscillations associated with choice expectations during choice. **a-b**. Time-frequency regression with Q-values after the cue onset and before the response respectively. Yellow colors indicate positive significance (cluster-corrected, p_c_ < 0.05). The horizontal dashed line represents the boundaries of the explored 4-12 Hz low-frequency oscillations range. **c-d**. Time-course of regression estimates with Q-values in the 4-12 Hz frequency range after the cue onset and before the response respectively. Average regression estimates±SEM (represented by a shaded gray area around the mean) across recording sites (n = 48 sites) plotted separately in the reward (Qr, green) and punishment (Qp, red) conditions. Colored horizontal bars indicate significant clusters (cluster-corrected, p_c_ < 0.05) in the time domain for a one-sample t-test against 0 in the reward (green) and punishment conditions (red). Black horizontal bars indicate the significant cluster (cluster-corrected, p_c_ < 0.05) in the time domain for the paired t-test comparing the regression estimates in the reward and punishment conditions. Reaction times (RT) in the reward and punishment conditions are represented as circles (reward: green; punishment: red) and horizontal lines (mean±SEM).

At the time of outcome display, we found that LFOs were positively associated with expectations (Fig. 3a) and negatively associated with the magnitude of the outcome (Fig. 3b). This demonstrates that the two core components of the teaching signal - the prediction-error - are encoded by thalamic LFOs which relate to the difference between what subjects expect and the actual decision outcome – what we get. Interestingly, around outcome onset, the level of expectation was significantly related to LFOs only in the reward-based learning condition (Fig. 3c; −0.66 to 1.06 s window, β_Rr_ = 0.17±0.01, sum(t(47)) = 75.92, p_c_ < 0.05), while outcomes were significantly encoded by LFOs in both rewarding and punishing conditions (Fig. 3d; 0.28 to 2.08 s window, β_Rr_ = −0.16±0.01, sum(t(47)) = −107.65, p_c_ < 0.05; 0.13 to 2.78 s window, β_Rp_ = −0.16±0.01, sum(t(47)) = 153.31, p_c_ < 0.05). Altogether, outcome-related activity is consistent with a similar encoding of rewards and punishments in the thalamus. Q-value encoding was detected only in the reward condition, but the absence of a significant difference between the two conditions prevent a conclusion in favor of a proper dissociation in the encoding of the prediction error (Supplementary Fig. S2).

**Figure 3.**
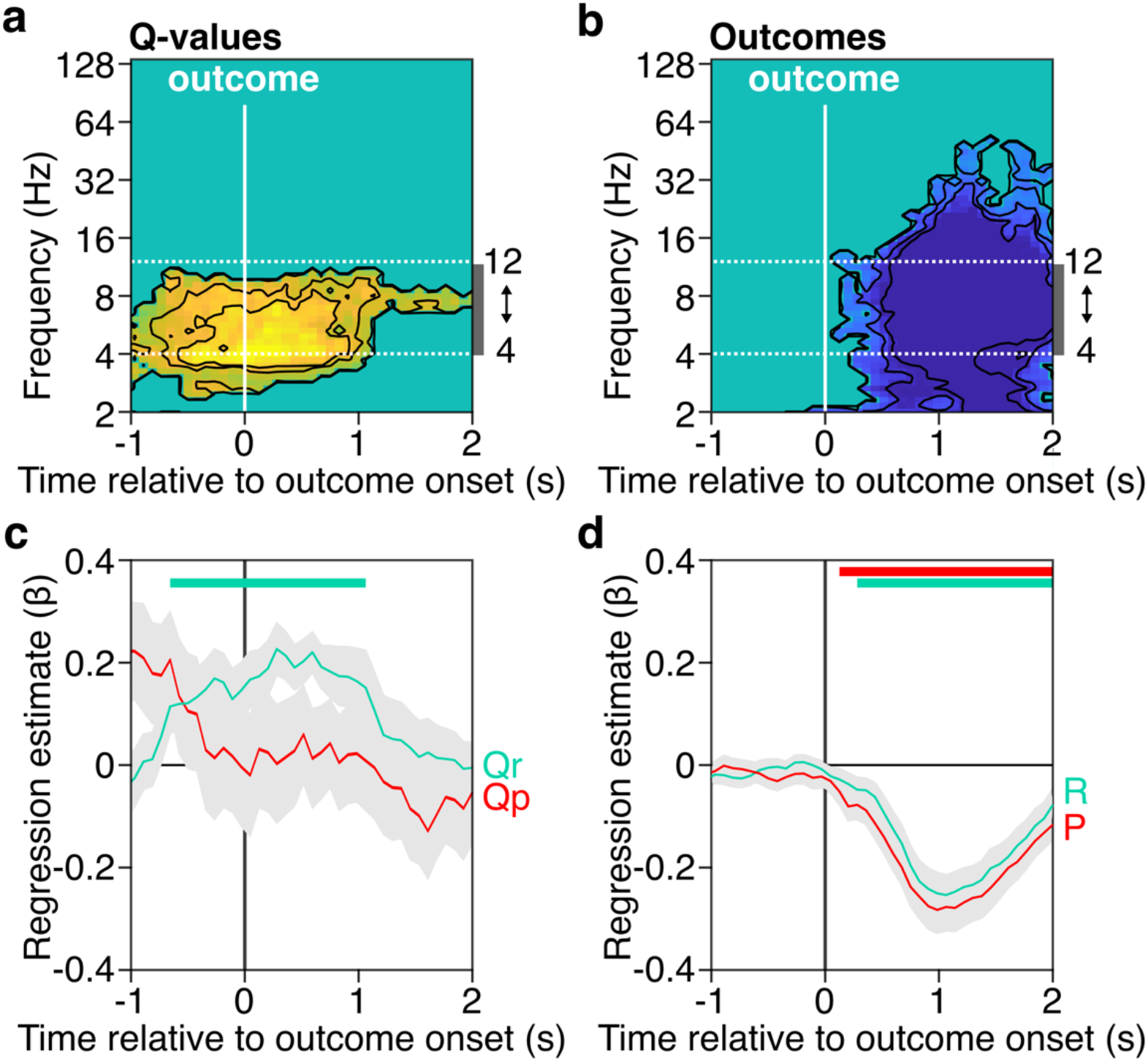
Thalamic low-frequency oscillations associated with prediction error components. **a-b**. Time-frequency decomposition of prediction error signals, with regression with Q-values and outcome values respectively. Blue and yellow colors indicate respectively negative and positive significance (cluster-corrected, p_c_ < 0.05). The horizontal dashed line represents the boundaries of the explored 4-12 Hz low-frequency oscillations range. **c-d**. Time-course decomposition of PE signals in the 4-12 Hz frequency range, with regression with Q-values and outcome values respectively. Average regression estimates±SEM (represented by a shaded gray area around the mean) across recording sites (n = 48) plotted separately in the reward (Qr/R, green) and punishment (Qp/P, red) conditions. Colored horizontal bars indicate significant clusters (cluster-corrected, p_c_ < 0.05) in the time domain for a one-sample t-test against 0 in the reward (green) and punishment conditions (red). No significant cluster (cluster-corrected, p_c_ < 0.05) in the time domain was found for the paired t-test comparing the regression estimates in the reward and punishment conditions.

Combining intra-thalamic human recordings with a probabilistic reinforcement learning task and trial-wise estimates of prediction errors from a Q-learning model brings the first mechanistic understanding of the role of the human limbic thalamus during reward-based vs. punishment avoidance learning. We found that during the choice phase, LFOs were better associated with punishment expectation signals, extending the previously observed role of the limbic thalamus in memory encoding in humans^22^ to aversive contexts which were examined in rabbits in early studies^15^. These signals could originate from the anterior insular cortex which was previously shown to implement punishment avoidance signals during an identical task in previous neuroimaging^11^ and intracranial^12^ studies.

Given the behavioral asymmetry in decision times between reward and punishment-based learning, we hypothesized that the neural activity could reflect the activation/inhibition balance of the thalamocortical learning circuitry during choice: the motor action threshold. This interpretation is also consistent with the fact that the Pavlovian bias on reaction times has been computationally interpreted as being largely due to an increase of non-decision time, which, within the decision diffusion modeling framework, is the parameter that better captures motor inhibition^7,23^.

Conversely, at the time of outcome processing, thalamic LFOs clearly encoded reward prediction errors. This likely reflects a cortical input from the ventromedial prefrontal cortex / lateral orbitofrontal cortex which was previously demonstrated to exhibit the same signals^12^. This finding echoes recent studies in non-human primates suggesting that LFOs oscillations in the orbitofrontal cortex are crucial for reward-guided learning and are driven by LFOs in the hippocampus^24^. As the limbic thalamus shares extensive connections with the hippocampus, orbitofrontal, and prefrontal areas, they may form together a circuit in which reward-guided learning is encoded by LFOs. Evidence for punishment prediction errors encoding in the thalamus was somehow weaker, if not incomplete. If confirmed, these results could be easily accommodated by the fact that several other brain areas and systems outside the fronto-striatothalamic circuits and devoted to punishment avoidance learning^11,12,13,14^.

Our results also allowed us to address another open question in the field, which is to test the frequency bands involved during learning. In mice, beta (13-30 Hz) synchrony between the mediodorsal thalamus and the prefrontal cortex was associated with learning^16^, whereas in humans, intracranial recording revealed that broadband gamma activity (50-150 Hz) recorded in the cortex encoded reward and punishment-based learning signals^12^.

To conclude, our study represents a step forward in elucidating the computational decisionmaking processes underlain by the thalamus. Given the centrality of this brain structure within the fronto-striatal circuit, we believe that understanding its function will prove useful to computationally characterize cognitive deficits observed in many neuropsychiatric disorders^25^.

## Methods

### Patients and surgical approach

Intracerebral recordings were obtained from 8 patients (38.1±3.7 years old, 3 females, see demographical details in Table S1) suffering from intractable epilepsy. They were implanted bilaterally in the limbic thalamic nuclei within the anterior thalamic nuclei (ATN) with deep-brain stimulation electrodes (Medtronic DBS lead model 3389, 4 contacts, 1.5 mm wide with0.5 mm spacing edge to edge between contacts) as a surgical treatment to alleviate their seizures. The stereotaxic trajectory of the electrode was calculated pre-operatively based on the patient’s MRI images. Electrodes were implanted through the ATN to ensure its maximal recording, with at least the two most dorsal contacts inside the ATN. As a result, the more ventral-proximal contacts pointed towards the dorsomedial thalamic nuclei (DMTN) located below the ANT along the implantation trajectory. Electrode implantation was performed according to the clinical procedures of the clinical trial “France” (NCT02076698), with targeted structures preoperatively selected according strictly to clinical considerations with no reference to the current study. Patients were investigated either in the epilepsy departments of Grenoble, Paris, or Marseille. All participants gave written informed consent and the study received approval from the ethics committee (Comité de Protection des Personnes Sud-Est I, protocol number: 2011-A00083-38).

### Behavioral task

Patients performed a probabilistic instrumental learning task. No seizures took place during the testing sessions. Patients were provided with written instructions (reformulated orally if necessary) stating that the goal was to maximize their financial payoff by considering rewardseeking and punishment avoidance as equally important. Patients performed short training sessions to familiarize themselves with the timing of events and with response buttons. Participants performed up to 6 sessions (see supplementary table 1). Each session was an independent task containing four new pairs of cues to be learned, each pair of cues being presented 24 times for a total of 96 trials. Cues were abstract visual stimuli taken from the Agathodaimon alphabet. The four cue pairs were divided into two conditions (2 pairs of reward and 2 pairs of punishment cues), associated with different pairs of outcomes (winning 1€ versus nothing or losing 1€ versus nothing). To win money, patients had to learn by trial and error the cue-outcome associations and choose the most rewarding cue in the reward condition and the less punishing cue in the punishment condition. The reward and punishment conditions were intermingled in a learning session and the two cues of a pair were always presented together. Within each pair, the two cues were associated with the two possible outcomes with reciprocal probabilities (0.75/0.25 and 0.25/0.75). On each trial, one pair was randomly presented, and the two cues were displayed on the left and right of a central fixation cross, their relative position being counterbalanced across trials. The subject was required to choose the left or right cue by using their left or right index to press the corresponding button on a joystick (Logitech Dual Action). Since the position on the screen was counterbalanced, response (left versus right) and value (good versus bad cue) were orthogonal. The chosen cue was colored in red for 250 ms and then the outcome was displayed on the screen after 1000 ms. Visual stimuli were delivered on a 19-inch TFT monitor with a refresh rate of 60 Hz, controlled by a PC with Presentation 16.5 (Neurobehavioral Systems, Albany, CA).

### Local field potentials acquisition and processing

Intracranial signals recordings were performed at the bedside of patients from externalized electrode leads in the two days following electrode implantation (i.e., before the electrodes were connected to the stimulator). Local field potentials were recorded from a bipolar montage between adjacent electrode contacts. Data were bandpass filtered online from 0.1 to 200 Hz and recorded either at 1024 Hz or 2048 Hz. Each electrode trace was subsequently re-referenced with respect to its direct neighbor (bipolar derivations with a spatial resolution of 2 mm) to achieve high local specificity by canceling out effects of distant sources that spread equally to both adjacent contacts through volume conduction. Overall, 48 sites were recorded (3 contact pairs/electrode × 2 hemispheres × 8 patients) using a commercial video-EEG monitoring system (System Plus, Micromed).

Time-frequency analyses were performed with the FieldTrip toolbox for MATLAB. The electrophysiological data were resampled at 512 Hz and segmented into epochs from 4s before to 4s after the cue onset and outcome onset. A multi-tapered time-frequency transform allowed the estimation of spectral powers (Slepian tapers; lower-frequency range: 1–32 Hz, 6 cycles and 3 tapers per window; higher frequency range: 32–200 Hz, fixed time-windows of 200 ms, 4–31 tapers per window). This approach uses a steady number of cycles across frequencies up to 32 Hz (time window durations, therefore, decrease as frequency increases) whereas, for frequencies above 32 Hz, the time window duration is fixed with an increasing number of tapers to increase the precision of power estimation by increasing smoothing at higher frequencies. Time-frequency power was converted into dB (decimal logarithm transformation) and z-scored to improve the Gaussian distribution of the data.

### Behavioral analysis and modeling

The percentage of correct choice (i.e., selection of the most rewarding or the less punishing cue) and reaction time (between cue onset and choice) were used as dependent behavioral variables. Statistical comparisons between the correct choice rate and chance choice rate (i.e., 0.5) were assessed using t-tests. Statistical comparisons of correct choice rate and reaction times between reward and punishment conditions were assessed using paired t-tests.

A standard Q-learning algorithm (QL) was used to model choice behavior. For each pair of cues, A/B, the model estimates the expected value of choosing A (Qa) or B (Qb), according to previous choices and outcomes. The initially expected values of all cues were set at 0, which corresponded to the average of all possible outcome values. After each trial (t), the expected value of the chosen stimuli (say A) was updated according to the rule:

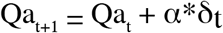

The outcome prediction error, δ(t), is the difference between obtained and expected outcome values:

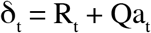

with R(t) the reinforcement value among −1€, 0€, and +1€. Using the expected values associated with the two possible cues, the probability (or likelihood) of each choice was estimated using the SoftMax rule:

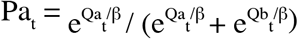

The constant parameters α and β are the learning rate and choice temperature, respectively. Expected values, outcomes, and prediction errors for each patient were then z-scored across trials and used as statistical regressors for electrophysiological data analysis.

### Regression between electrophysiological signals with reward and punishment learning behaviors

Power (Y) at each time-frequency point was regressed using a general linear model against both outcome value (R) and expected value (Q) to obtain a regression estimate for each timefrequency point and each contact pair:

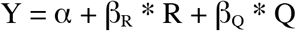

with β_R_ and β_Q_ corresponding to the R and Q regression estimates, respectively. The significance of regression estimates was assessed at each time-frequency point using a using one-sample two-tailed t-test against 0 across all sites. Permutation tests were performed to control for multiple comparisons. The pairing between power and regressor values across trials was shuffled randomly 60,000 times. The maximal cluster-level statistics (the sum of t-values across contiguous time points passing a significance threshold of 0.05) were extracted for each shuffle to compute a ‘null’ distribution of effect size. For each significant cluster in the original (non-shuffled) data, we computed the proportion of clusters with higher statistics in the null distribution, which is reported as the ‘cluster-level corrected’ p_c_ value. Low-frequency (4-12 Hz) time series were computed, and the same general linear model approach was used for each time point of the time series separately in the reward and punishment conditions. The significance of regressors was assessed using a cluster correction approach comparable to the one described above.

