## Supplementary Figures S1, S2, Supplementary Table 1 for "The role of the thalamus in human reinforcement learning"

### Supplementary materials

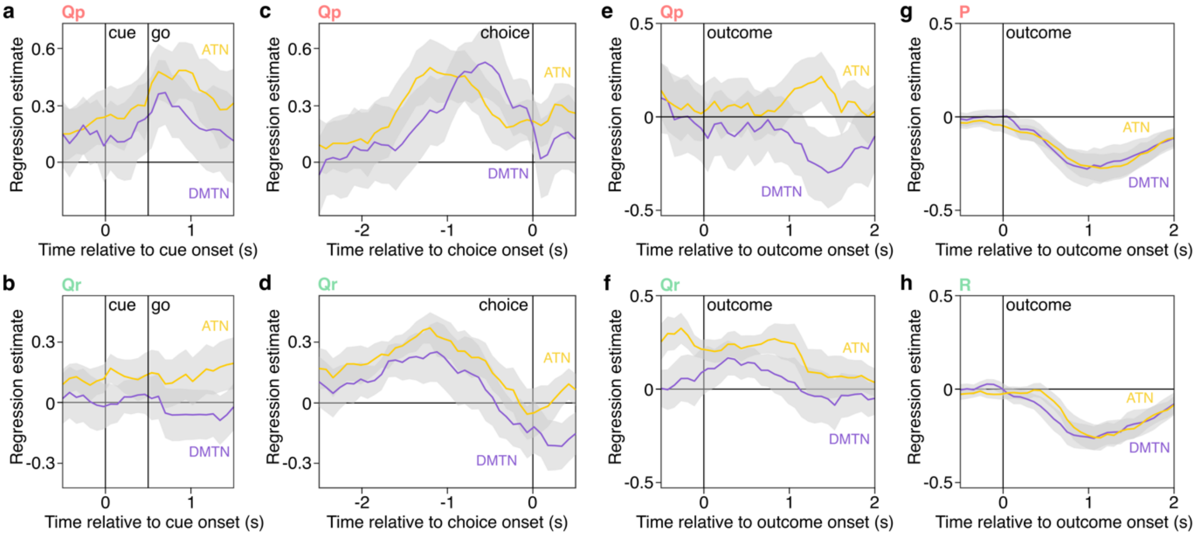


**Supplementary Figure S1. Comparison of low-frequency encoding of expectation and outcome between ATN and DMTN**

Time-course regression in the 4-12 Hz frequency range with Q-values (Qr, Qp) and outcome (R, P) values with regression estimate averaged±SEM (the shaded gray area around the mean) across recording sites plotted separately for ATN (yellow, n = 16 sites) and DMTN (purple, n = 16 sites) in the punishment (a, c, e, g) and reward (b, d, f, h) conditions. No significant cluster (cluster-corrected, $p_c < 0.05$) in the time domain was found for the paired t-test comparing the regression estimates in the ATN and DMTN.

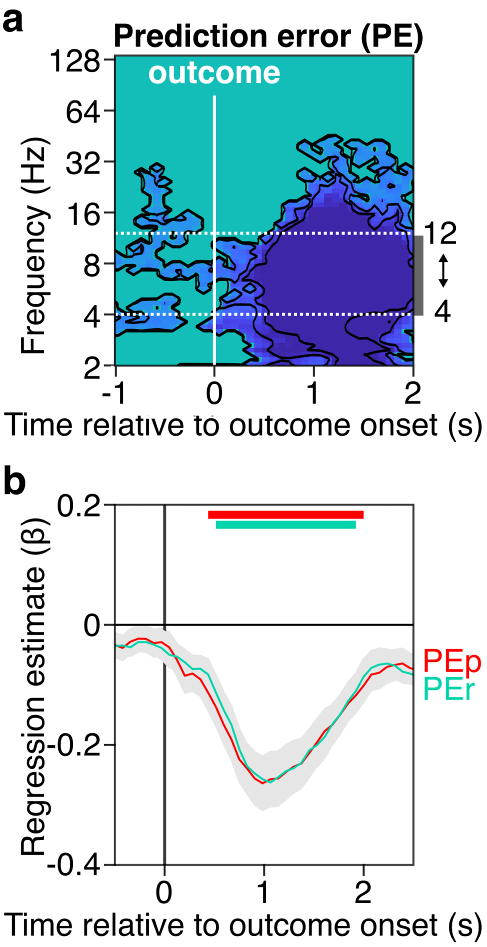


**Supplementary Figure S2. Thalamic low-frequency oscillations associated with prediction error**

**a-b.** Time-frequency regression with prediction error (PE). Blue colors indicate negative significance (cluster-corrected, $p_c$<0.05). The horizontal dashed line represents the boundaries of the explored 4-12 Hz low-frequency oscillations range. **c-d.** Time-course of PE regression estimates in the 4-12 Hz frequency range. Average regression estimates±SEM (represented by a shaded gray area around the mean) across recording sites (n = 48) plotted separately in the reward (PEr, green) and punishment (PEr, red) conditions. Colored horizontal bars indicate significant clusters (0.52 to 1.92 s window, $\beta_{PEr}$ = -0.20±0.01, sum(t(47)) = -110.11, $p_c$ < 0.001; 0.44 to 2.00 s window, $\beta_{PEp}$ = -0.19±0.01, sum(t(47)) = -108.09, $p_c$ < 0.001) in the time domain for a one-sample t-test against 0 in the reward (green) and punishment conditions (red). No significant cluster (cluster-corrected, $p_c$ < 0.05) in the time domain was found for the paired t-test comparing the regression estimates in the reward and punishment conditions.

| Inclusion center | Gender | Age (years) | Age at first seizure (years) | Epileptic syndrome | Seizure origin | Medication at surgery | Number of sessions of 96 trials |
|---|---|---|---|---|---|---|---|
| Grenoble | M | 29 | <1 | Frontal lobe epilepsy | bilateral | OXC, LAC, CLB, INN | 6 |
| Grenoble | M | 54 | 27 | Frontal lobe epilepsy | bilateral | LAC, CBZ, PRG | 6 |
| Grenoble | F | 35 | 5 | Temporo-occipital lobe epilepsy | bilateral | CLB, LZP, FBM | 6 |
| Grenoble | F | 34 | <1 | Hypothalamic hamartoma | bilateral | LAC, RFN, CLB | 6 |
| Grenoble | M | 25 | 7 | Parietal lobe epilepsy | right | FBM, LEV, VGT, CLN | 5 |
| Grenoble | F | 51 | 12 | Occipital lobe epilepsy | bilateral | VPA, LTG, CLB | 6 |
| Marseille | M | 44 | 32 | Temporo-frontal lobe epilepsy | bilateral | CBZ, LAC, LEV, PGB, CLB | 5 |
| Marseille | M | 33 | 7 | Multifocal epilepsy | bilateral | OXC, TPM, CLB | 4 |

**Table S1. Cohort data**

OXC: oxcarbazepine; LAC: lacosamide; CLN: clonazepam; CLB: clobazam; INN: retigabine; CBZ: carbamazepine; PRG: pregabaline; LZP: lorapezam; FBM: felbamate; RFN: rufinamide; LEV: levetiracetam; VGT: vigabatrin; VPA: valproic acid; LTG: lamotrigine; PGB: pregabaline; TPM: topiramate
